# Leukemia circulation kinetics revealed through blood exchange method

**DOI:** 10.1101/2023.09.03.556043

**Authors:** Alex B. Miller, Adam Langenbucher, Felicia H. Rodriguez, Lin Lin, Christina Bray, Sarah Duquette, Ye Zhang, Dan Goulet, Andrew A. Lane, David M. Weinstock, Michael T. Hemann, Scott R. Manalis

## Abstract

Leukemias and their bone marrow microenvironment are known to undergo dynamic changes over the course of disease. However, relatively little is known about the circulation kinetics of leukemia cells, nor the impact of specific factors on the clearance of circulating leukemia cells (CLCs) from the blood. To gain a basic understanding of leukemia cell dynamics over the course of disease progression and therapeutic response, we apply a blood exchange method to mouse models of acute leukemia. We find that CLCs circulate in the blood for 1-2 orders of magnitude longer than solid tumor circulating tumor cells. We further observe that: i) leukemia presence in the marrow can limit the clearance of CLCs in a model of acute lymphocytic leukemia (ALL), and ii) CLCs in a model of relapsed acute myeloid leukemia (AML) can clear faster than their untreated counterparts. Our approach can also directly quantify the impact of microenvironmental factors on CLC clearance properties. For example, data from two leukemia models suggest that E-selectin, a vascular adhesion molecule, alters CLC clearance. Our research highlights that clearance rates of CLCs can vary in response to tumor and treatment status and provides a strategy for identifying basic processes and factors that govern the kinetics of circulating cells.

## Introduction

Leukemias are hematopoietic cancers characterized by the transit of malignant cells through the blood^1^. They arise from mutations in cells within specific sites of hematopoietic development, but quickly spread via circulating leukemia cells (CLCs) to sites throughout the body and have heavy involvement of not only hematological organs such as lymph nodes and spleen, but also the liver and central nervous system ^2–4^. Several historical studies have examined the clearance rates of solid tumor circulating tumor cells (CTCs) in mice ^5–8^, which are the metastatic analogs of CLCs in nonhematological tumors^9,10^. However, these studies relied on collecting blood samples from different mice at discrete timepoints, or using continuous detection in small capillaries, which leads to low accuracy measurements due to mouse-to-mouse heterogeneity or under-sampling of events. Our group has developed a system for continuous monitoring of fluorescent CTCs through the carotid artery of mice, allowing for highly accurate measures of real-time CTC concentration^11^. We previously used this system to identify kinetics of solid tumor CTCs in mice, and reported that half-life times across a variety of tumor types range between 1 and 4 minutes^12^. However, the measurement of circulation kinetics of leukemia CLCs and comparison to solid tumor CTCs has not yet been explored. These measurements may help explain fundamental questions related to the biology of leukemias. For example, the higher burden of some leukemias observed in the blood could result from a greater rate of intravasation (*i*.*e*., more entry), lower rate of extravasation (*i*.*e*., less exit) or both.

In the context of active leukemia, the bone marrow niche undergoes a number of physical and functional changes. Compared to healthy marrow, leukemic marrow commonly exhibits hypercellularity, enhanced extracellular matrix deposition, vascular adhesion protein expression, and vascular permeability^13–17^. However, little is known regarding the circulation properties of leukemia cells or the factors that regulate circulation. Recent studies have suggested that adhesion proteins, including collagen, fibronectin, and selectins, play an important role in developing a protective niche within the bone marrow. E-selectin can support chemoresistance in acute myeloid leukemia (AML) through activation of pro-survival signaling, and integrin β1 adhesion can induce chemoprotective drug efflux in acute lymphocytic leukemia (ALL)^18–20^. Interfering with these adhesion pathways has been shown to increase chemotherapeutic efficacy and promote malignant cell egress from the bone marrow into the blood compartment^21–24^.

Additional studies have suggested that chemotherapeutic treatment of leukemias alters their transcriptional and phenotypic states compared to their untreated counterparts, including the upregulation of adhesion molecules^22,25^. Indeed, there is now extensive literature suggesting that elevated cell adhesion can mediate leukemia resistance to chemotherapy^26,27^. However, few studies have explored changes in leukemia cell circulation properties following disease relapse, and there has, consequently, been little functional validation that upregulation of adhesion molecules alters leukemia circulation kinetics.

Here, we address these questions by employing a blood exchange system to quantify and model circulation kinetics of both acute lymphocytic leukemia (ALL) and acute myeloid leukemia (AML) models. These data reveal that the half-life time of leukemia cells is over 40 times longer than that of solid tumor CTCs. We then vary the donor-recipient pairs to identify factors that govern the circulation. We find that the presence of ALL in the bone marrow at least 8 days post tumor initiation can limit the clearance of cells from the blood and can be reversed by blocking the vascular adhesion protein E-selectin. Finally, we show that CLCs from a relapsed AML model, but not an ALL model, have faster rates of clearance compared to an untreated version of the same model. This enhanced clearance rate is associated with higher expression of E-selectin ligands. Interfering with E-selectin decreases the clearance of relapse cells. Altogether, this platform identifies critical differences in circulation profiles between solid tumor CTCs and CLCs, as well as between untreated and relapsed leukemia, and provides a method for identifying the impact of specific factors, such as adhesion proteins, on CLC clearance in the blood.

## Results

### Blood exchange for measuring circulation properties of CLCs

To measure the clearance profile of CLCs, we used a modified version of our previously published blood exchange method^11,12^. By connecting the circulation of a fluorescent-leukemia-bearing donor mouse to a non-leukemia-bearing “healthy” recipient mouse, naturally shed fluorescent CLCs were intravenously infused into a recipient animal for 30 minutes to 1 hour (Fig. 1a). After disconnecting the mice, we performed a 3-hour post-blood exchange scan of the recipient mouse using a microfluidic fluorescent-detection platform to obtain real-time concentrations of CLCs in the blood. An exponential decay curve of the fluorescent CLCs was then fit to the plot of CLC concentration over time to estimate the clearance profile of CLCs in circulation. Two key properties of clearance were extracted from the decay profiles: fraction remaining and equilibration time. The fraction remaining describes the relative change in concentration over the course of the 3-hour scan and estimates what fraction of the cells are still in circulation by the end of that time period (whether or not they exited and then reentered circulation), assuming that our continuous detection of fluorescent CLCs in the carotid artery is representative of the overall CLC concentration in the mouse’s blood. It can be calculated by comparing the CLC concentration at the beginning and end of the post-blood exchange scan using the following equation:

**Figure 1.**
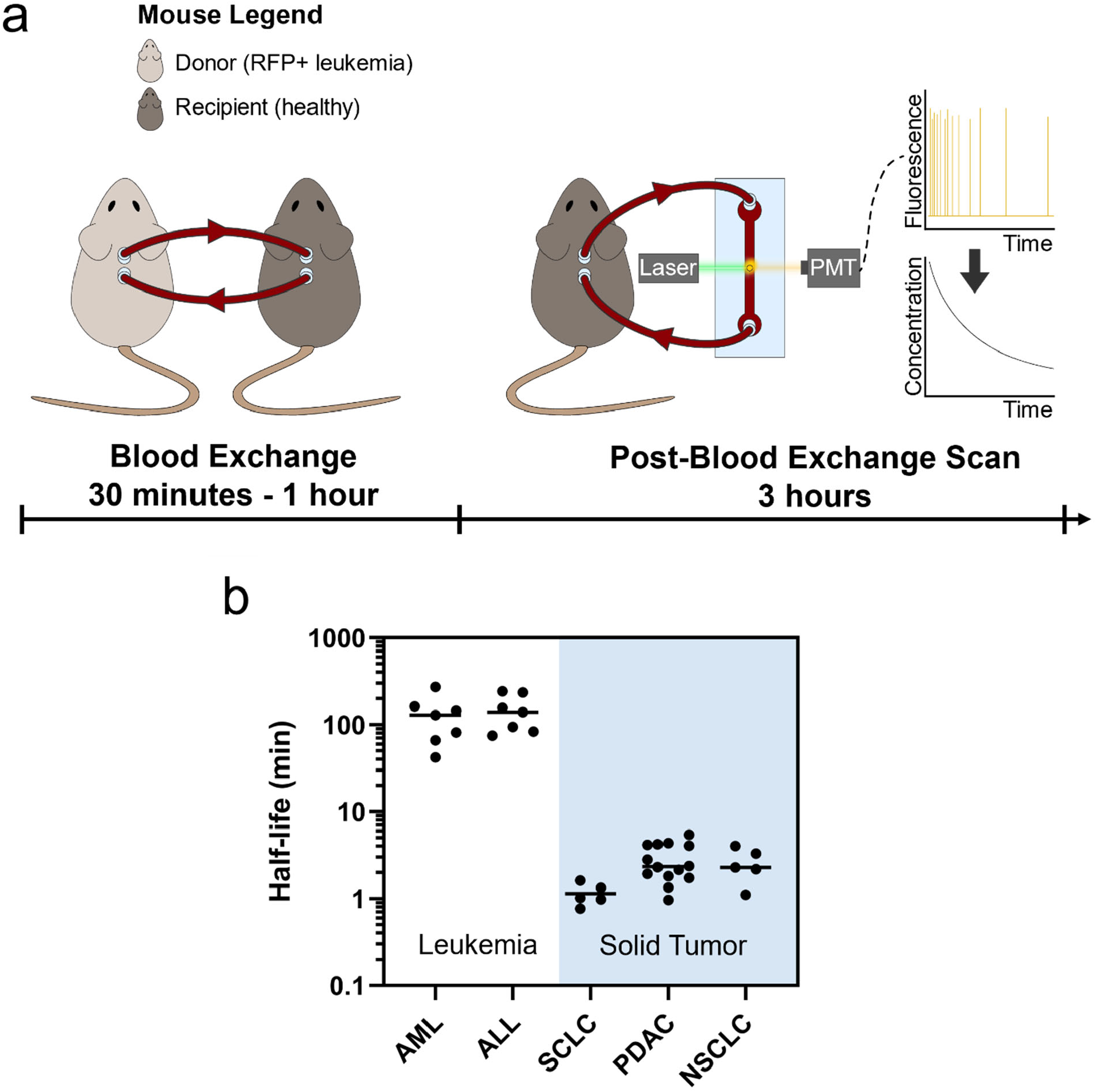
Blood exchange identifies differences in clearance kinetics of circulating tumor cells between leukemia and solid tumor models. (a) Overview schematic of blood exchange setup. A donor and recipient mouse have their circulation connected for 30 minutes to 1 hour. Following this blood exchange period, the mice are disconnected and the circulation from the recipient mouse is monitored for three hours. The blood passes through a laser on a microfluidic chip, and emitted light from fluorescent CLCs is detected by a photomultiplier tube (PMT). The raw data from the PMT is then processed to identify the concentration of CLCs, and the decay profile is used to extract circulation parameters. (b) Half-life times of circulating leukemia cells (AML and ALL) is several orders of magnitude higher than historical measures of various solid tumor circulating tumor cells^12^. SCLC: small-cell lung cancer; PDAC: pancreatic ductal adenocarcinoma; NSCLC: non-small-cell lung cancer. Each dot represents an independent donor-recipient mouse pair.

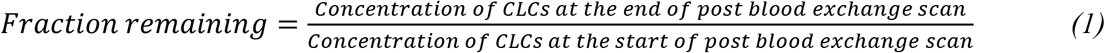

The second parameter used to characterize the decay profile was the equilibration time. This parameter is similar to a half-life time and describes how long it takes for the concentration in the blood to reach a steady state. Since not all of the curves decay to zero, it was important to add a constant to the equation, such that there was a decay to constant, rather than a decay to zero (a decay to zero describes a half-life measurement). A best fit curve was applied to the decay data, with inputs of concentration and time in minutes, using the following equation:

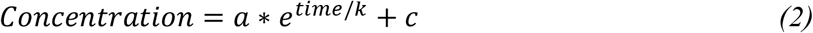

where k is the equilibration time in minutes. The parameters a, k, and c were swept over positive values to find the best fit and identify the estimated equilibration time.

We started by testing two fluorescent syngeneic models of leukemia to study circulation kinetics, a BCR-ABL driven B cell ALL model and an MLL-AF9 driven AML model^28–33^, each paired with healthy recipients. In these experiments, the calculated constant “c” in the exponential decay equation was found to be near 0, so the equilibration times in these experiments are equivalent to the true half-lives of the CLCs in circulation. Although the circulating concentration of CLCs varied by at least 1000-fold over the 3-week time course of disease (Supp. 1a), we observed no significant correlation between days post leukemia initiation and either half-life or fraction remaining (Supp. 1b,c), suggesting that the early and later stage CLCs have similar circulation properties in both models.

In a previous study we measured the half-life times of CTCs from three solid tumor models: small cell lung cancer (SCLC), pancreatic ductal adenocarcinoma (PDAC) and non-small cell lung cancer (NSCLC)^12^. Since the solid tumor half-life estimates were also performed through blood exchange with healthy recipient mice, we could compare the half-life times in circulation of the solid tumor CTCs to the CLCs from the leukemia models. Interestingly, we found the half-life times of CLCs to be approximately 200 minutes, compared to between 1 and 5 minutes for the solid tumor models (Fig. 1b). This finding demonstrates that, among the models we tested, solid tumor CTCs have markedly faster clearance in the blood compared to CLCs.

### Impact of bone marrow cellularity on clearance of CLCs

A long-standing question in both the leukemia and transplantation fields is whether bone marrow occupancy impacts the dissemination of cells introduced into the circulation^34^. To assess whether the presence of leukemia in the bone marrow can impact the clearance rates of CLCs, we varied the leukemia status of recipient animals in our blood exchange clearance experiments (Fig. 2a). We used a non-fluorescent (RFP-) version of the same BCR-ABL driven ALL leukemia tested previously. We started by comparing clearance rates when RFP+ ALL donor mice were connected to healthy recipient mice or to recipients bearing active RFP-ALL with at least 30% leukemia burden in the marrow. We found a striking difference in the clearance profiles (Fig. 2b,c: “Healthy Mouse” and “Leukemia”). While clearance of RFP+ ALL CLCs followed a steady decrease over the 3-hour post-blood exchange scan when infused into healthy recipients, there was an 84% increase in the fraction remaining term (Equation 1) with RFP-ALL-bearing recipients.

**Figure 2.**
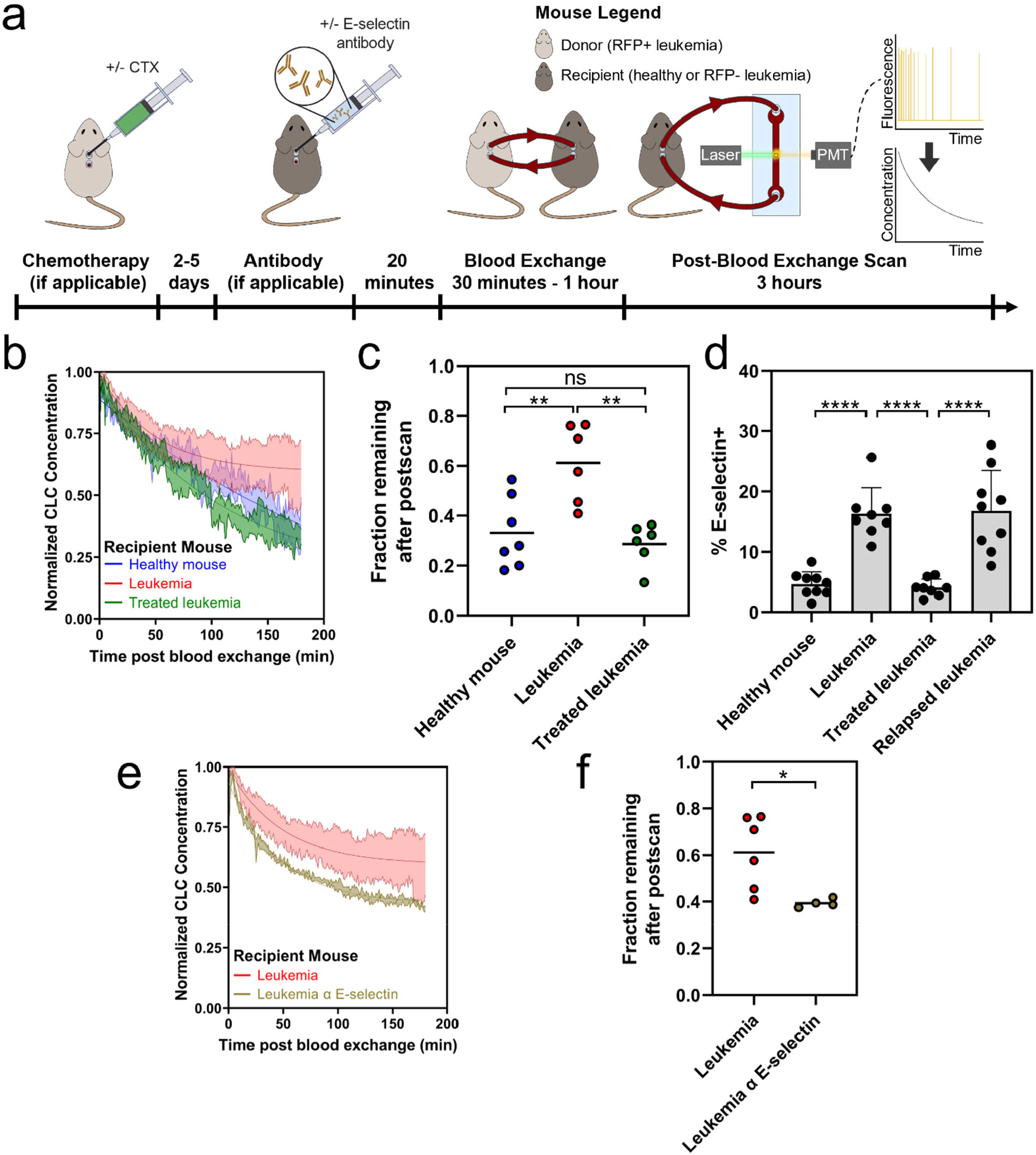
Bone marrow microenvironment can impact clearance of CLCs. (a) Overview of experimental plan. Some mice (panels e and f) are dosed with E-selectin antibody prior to blood exchange and postscan (b) Decay profiles of RFP+ CLCs in recipients with varied tumor status. (c) Decreased clearance of CLCs in recipient mice with active disease as measured by an increase in fraction of CLCs remaining in circulation (Tukey multiple comparisons **p<0.01). (d) E-selectin expression on BMECs increases in diseased context (bars represent mean +/-standard deviation; Tukey multiple comparisons ****p<0.0001). (e) Decay profiles of RFP+ CLCs in tumor bearing recipient mice with or without E-selectin antibody treatment (f) Dosing recipient mice with E-selectin antibody allows for increased clearance of CLCs in diseased recipients as measured by decrease in fraction remaining (two-tailed t test p = 0.0254). For decay profiles (b and e), shaded regions are represented by mean +/-standard error, and lines represent the best fit decay curve. For fraction remaining analysis (c and f), each dot represents an independent donor-recipient mouse pair.

It is worth noting that young adult mouse bone marrow is highly cellular (typically >95%) so the longer equilibration time in leukemic recipients is less likely to simply result from less “availability of space” for engraftment and more likely result from remodeling induced by the engrafted leukemia. To test this hypothesis, we treated mice engrafted with RFP-ALL with a one-time dose of cyclophosphamide (CTX) and performed blood exchange 2-5 days post treatment. This dose resulted in a decrease in percentage of leukemia cells to below 5% through day 5 post treatment, and approximately a 3-week survival extension (see Methods). In the CTX-treated mice, the clearance profile returned to that of the healthy mice (Fig. 2b,c: “Treated leukemia”). The data showed an approximately 2-fold increase in fraction remaining in the diseased (RFP-) recipients compared with healthy mouse (HM) recipients, about 0.6 compared to 0.3. This phenotype was fully reversed with CTX treatment, where the clearance of CLCs in recipient mice returned to a fraction remaining of approximately 0.3. These preliminary findings suggest that the presence of an existing leukemia burden actively reduces the extravasation of CLCs in a manner that rapidly reverses with treatment. Of note, because the changes were reversed within several days of CTX treatment, it is unlikely that such changes are due to vascular remodeling since these processes can take up to three weeks to be observed^35^.

As a next step, we further addressed whether the reduced bone marrow cellularity from treatment directly promotes CLC extravasation. To that end, we compared the clearance rates of RFP+ ALL CLCs in healthy recipients that had been treated either with CTX or irradiation. First, we confirmed that both chemotherapy treatment and irradiation induced hypocellularity in the bone marrow of healthy mice (Supp. 2a), and quantification revealed a drop in total number of cells in the marrow of approximately 50-75% (Supp. 2b). We then performed blood exchange experiments using ALL-bearing donor mice and monitored the decay in the post-blood exchange scan (Supp. 2c). While there were small changes in the clearance profiles, there were no significant decreases in the fraction remaining at the end of the post-blood exchange scan in either the irradiated or chemotherapy-treated recipients (Supp. 2d). These preliminary data suggest that engraftment kinetics may be affected by disease state more than bone marrow cellularity. This is noteworthy considering varied clinical approaches to recipient conditioning and bone marrow clearance prior to hematopoietic stem cell or T cell transplantation that could be studied using this approach^36,37^.

### Impact of bone marrow adhesion mediators on clearance of CLCs

To gauge the ability of this approach to monitor leukemia cell clearance kinetics following perturbations to the leukemia microenvironment, we focused on E-selectin, an endothelial surface protein. E-selectin attaches to several glycans and glycoproteins by binding to sialylated carbohydrates^38^, is important in leukocyte recruiting, and is upregulated in response to inflammatory cytokines, including TNF-α (tumor necrosis factor α) and IL-2 (interleukin 2)^1^. Additionally, a previous study showed that the inhibition of E-selectin led to an increase in the circulating number of CLCs in a model of AML^22^. To determine the potential relevance of E-selectin on our ALL model, we monitored E-selectin levels over the course of leukemia treatment. We also allowed the treated mice to relapse (>50% in the bone marrow at 2.5 weeks post treatment), and measured E-selectin expression at relapse. Notably, we found an increase in the percent of E-selectin positive bone marrow endothelial cells (BMECs) from about 5% for healthy and acutely treated mice to around 15% in diseased and relapse mice (Fig. 2d). This finding is similar to what we observed in the blood exchange experiments with diseased and treated mice, where a shift in leukemia clearance kinetics with a diseased recipient was reversed through treatment. Notably, a different endothelial binding protein, VCAM1, which binds to the integrin VLA4 (very late antigen 4 α4β1) and has been implicated in regulating ALL chemotherapy resistance^18,21^, was found to not correlate with disease state (Supp. 3). The observed increase of E-selectin on BMECs in the context of disease, and subsequent decrease upon treatment has not previously been explored in ALL but matches previously reported findings of acutely treated AML. Historical data suggests AML cells create inflammatory signaling that upregulates E-selectin expression on BMECs^22^, though no assessment of E-selectin expression on BMECs at late relapse has previously been reported. This association between leukemia status and E-selectin expression was also observed on BMECs from the AML model (Supp. 4).

Since E-selectin expression on BMECs changed over the course of disease and treatment, we decided to test whether inhibiting E-selectin using blocking antibodies would reverse the shift in clearance kinetics observed from blood exchange experiments. We used RFP+ ALL-bearing mice as the donors of blood exchange with RFP-ALL-bearing recipients treated with 100µg of α-E-selectin antibody or vehicle control 20 minutes before the start of blood exchange. Consistent with the previous study in AML^22^, we found that dosing with E-selectin antibody intravenously was able to increase the circulating concentration of ALL CLCs within 20 minutes (Supp. 5). The decay profile of RFP-mice dosed with E-selectin antibody showed a distinct shift from the non-dosed RFP-recipient mice (Fig. 2e), and the fraction remaining at the end of post-blood exchange scan was found to be significantly lower (p = 0.0254) in diseased mice treated with E-selectin antibody (Fig. 2f). These results show that disrupting E-selectin binding increases the number of ALL CLCs, at least in part, by reducing extravasation. Additionally, these data suggest that our approach can provide high resolution measurements of leukemia kinetics following perturbations that potentially impact leukemia cell retention in bone marrow or other sites of persistent disease.

### Impact of Relapse Status on Clearance of CLCs

To address whether cellular changes after chemotherapy can alter the circulation profile of CLCs, we used treated and untreated leukemia-bearing animals as donors in our blood exchange pairs. This allowed for the study of how untreated or relapsed CLCs clear in healthy recipient mice and served as a method for decoupling changes imparted from the bone marrow microenvironment with those imparted from the leukemia cells themselves. We performed these experiments using both the ALL and AML models. For these experiments, relapse was defined by leukemia regrowth to at least 50% in the marrow following treatment and nadir. In our ALL model, we saw similar clearance kinetics in the recipient healthy mice with either untreated or relapse donor CLCs, with no change in the equilibration time or fraction remaining (Fig. 3a,b, Supp. 6a). However, in the relapsed AML model, CLCs had a more rapid initial clearance (Fig. 3c,d, Supp. 6b). Quantitatively, this was reflected in the equilibration times, where relapse CLCs had a significantly faster equilibration time than the untreated CLCs, with over 100 minutes for untreated CLCs compared to around 20 minutes for the relapse CLCs.

**Figure 3.**
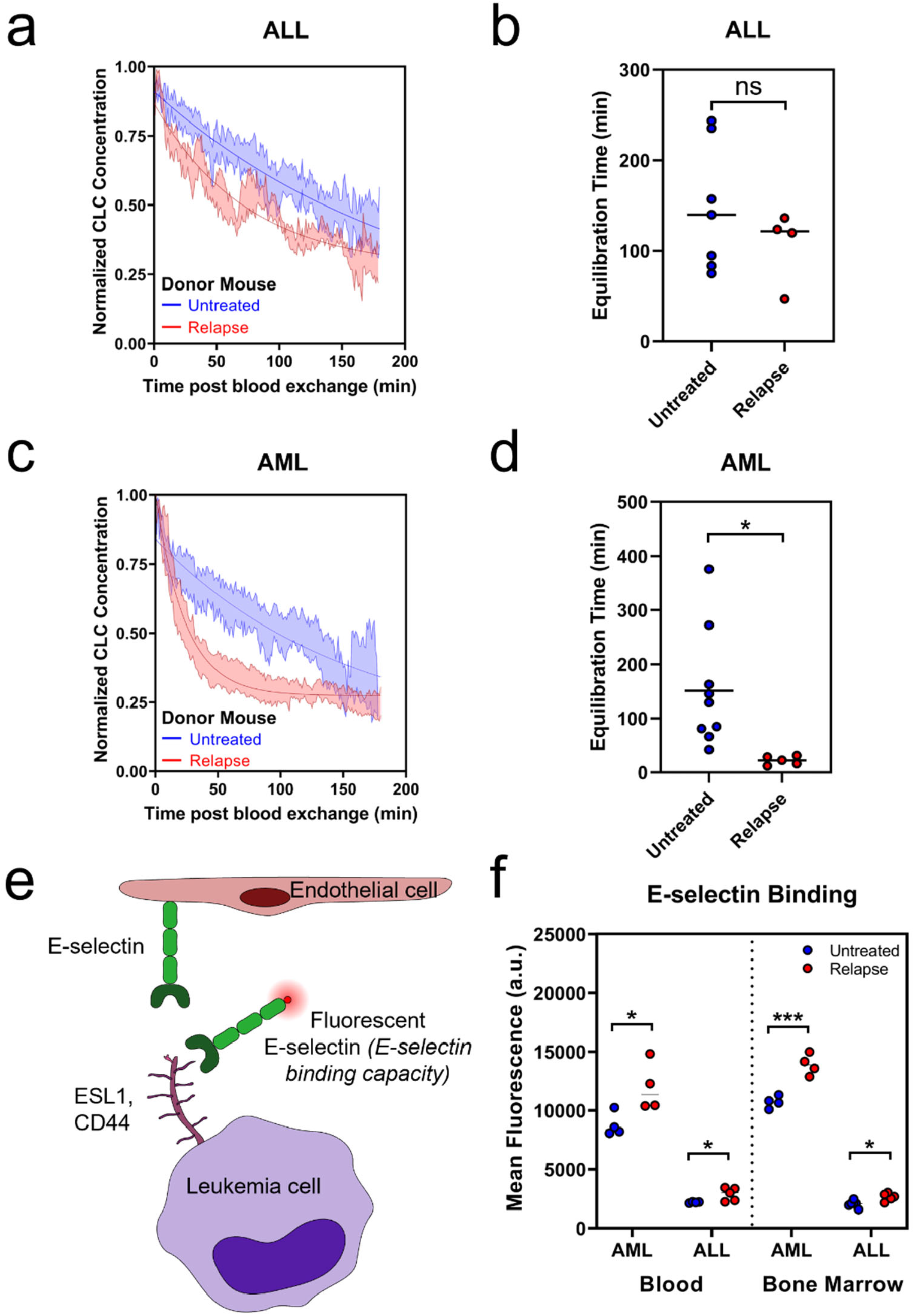
Blood exchange identifies differences in clearance kinetics in healthy mice with either untreated or relapse donors. (a) Decay profiles of untreated and relapse ALL CLCs in healthy recipient mice. (b) Relapse and untreated ALL CLCs have similar kinetics as measured by equilibration time (two-tailed t test p=0.3257) (c) Decay profiles of untreated and relapse AML CLCs in healthy recipient mice. (d) Equilibration time of relapse AML CLCs significantly faster than untreated (two-tailed t test p=0.0230). (e) Schematic of E-selectin adhesion to endothelial cells and E-selectin binding assay using fluorescent E-selectin. (f) E-selectin binding capacity increases in blood and marrow at relapse in AML and ALL (two-tailed t tests: AML Blood p=0.0330, ALL Blood p=0.0443, AML Bone Marrow p=0.0008, ALL Bone Marrow p=0.0112). For decay profiles (a and c), shaded regions are represented by mean +/-standard error, and lines represent the best fit decay curve. For equilibration time analysis (b and d), each dot represents an independent donor-recipient mouse pair.

We next wondered if changes in E-selectin could underlie the altered CLC kinetics between untreated and relapsed AML. Because E-selectin, found on endothelial cells, binds to sialyl groups of glycans and glycoproteins, there is no single antibody to effectively assess the ability of leukemia cells to bind E-selectin^39^. As such, we tested E-selectin binding capacity by incubating the leukemia cells with a fluorescently conjugated E-selectin protein (using a chimera of recombinant mouse E-selectin and human IgG with a fluorescent α-human IgG antibody) (Fig. 3e). Using flow cytometry, we compared the capacity to bind E-selectin between untreated and relapsed disease in both the blood and bone marrow (Fig. 3f). The binding capacity in the AML model at both untreated and relapse states was markedly higher than in the ALL model, and the increase in E-selectin binding at relapse was much greater in the AML model than the ALL model. Integrin β1, which has been suggested to contribute to chemoprotective niche, showed similar trends (Supp. 7). Our findings suggest that the slight increase in binding capacity observed in ALL cells may not be sufficient to alter the clearance rates, while the dramatic increase in E-selectin binding in AML can sufficiently alter the rate of clearance of CLCs.

The binding capacity of E-selectin and expression of integrin β1 increased acutely post chemotherapy in the bone marrow in both models and remained elevated compared to pre-treatment at relapse (Supp. 8). While other studies have similarly shown increases in adhesion expression directly following treatment^22^, there has previously been little evidence of whether that expression remains increased at relapse.

We have previously shown that buoyant mass can be a proxy for passage time through a channel^40^. Using a suspended microchannel resonator to measure the buoyant mass of single cells^41–43^, we found that there were no significant differences between untreated and relapsed disease for either model in either blood or bone marrow (Supp. 9). This suggests that difference in size does not explain the change in circulation that was observed.

Since E-selectin inhibition on endothelial cells was sufficient to alter clearance kinetics in a diseased ALL mouse (as described in Fig. 2), we wanted to explore whether interfering with E-selectin binding of leukemia cells could affect the observed changes associated with treatment in a different disease model. To that end, we dosed relapsed AML mice with recombinant mouse E-selectin 20 minutes prior to blood exchange with a healthy recipient, in order to occupy glycoproteins on the leukemia cell surface (Fig. 4a). We found that mice engrafted with relapsed AML dosed with recombinant E-selectin showed a significantly longer equilibration time and higher fraction remaining in circulation compared to non-treated relapse mice (Fig. 4b-d). These results suggest that the increase in E-selectin binding molecules on AML relapse cells increases the ability of the relapse cells to exit circulation faster and demonstrates that inhibiting E-selectin binding can prevent CLCs from rapidly leaving circulation.

**Figure 4.**
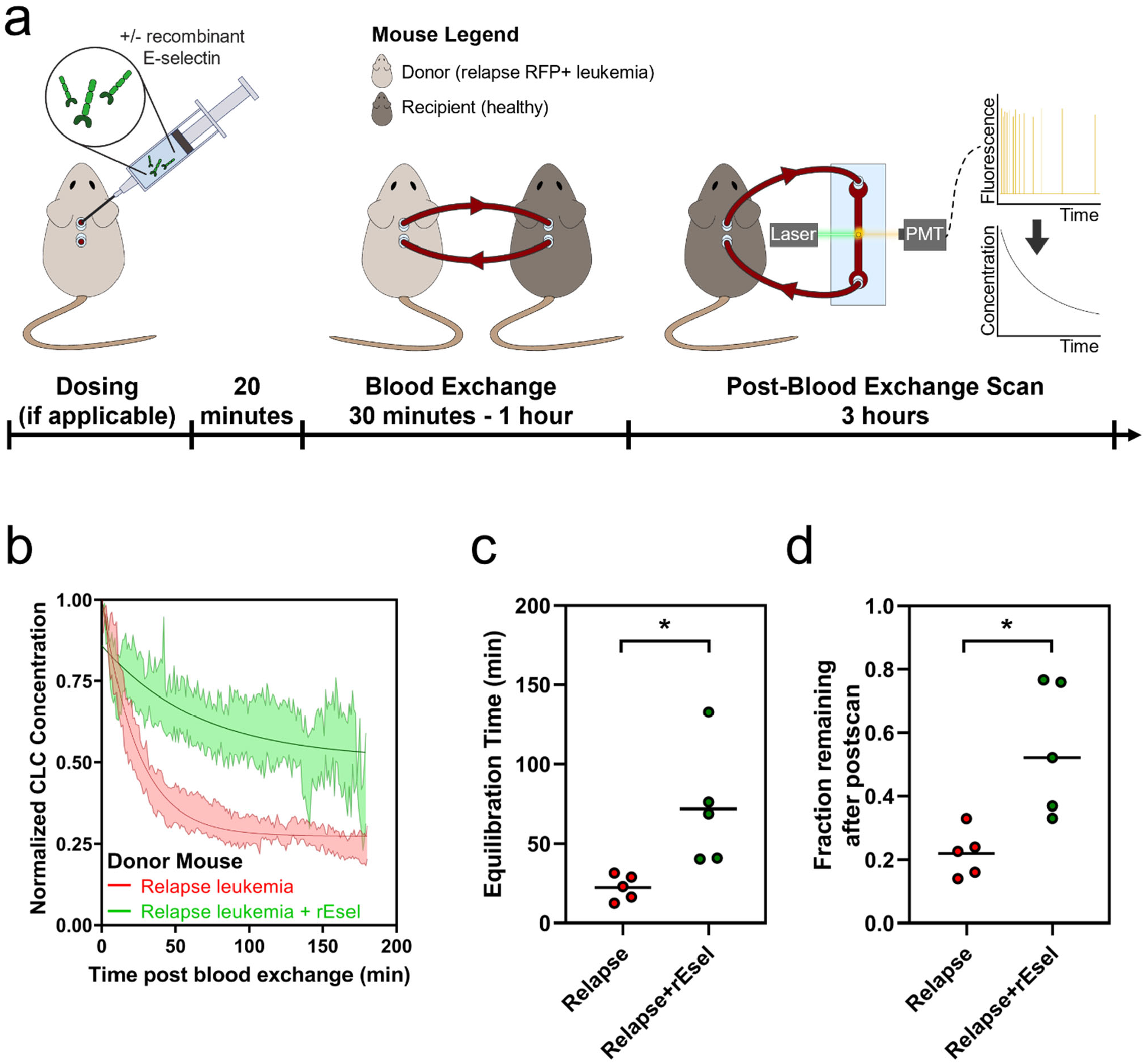
Interfering with E-selectin binding decreases clearance of relapse AML CLCs. (a) Overview of experiment. Recombinant E-selectin is added to the relapse donor mouse prior to blood exchange with healthy recipient. (b) Decay profiles of relapse CLCs with or without recombinant E-selectin (rEsel) treatment in healthy recipient mice (c, d) CLCs from relapse mice treated with recombinant E-selectin showed increase in both equilibration time (c, two-tailed t test p=0.021) and fraction remaining (d, two tailed t-test p=0.010). For decay profiles (b), shaded regions are represented by mean +/-standard error, and lines represent the best fit decay curve. Each dot represents an independent donor-recipient mouse pair.

## Discussion

The data presented here highlight the capability of the blood exchange system to quantify the impact of biological processes and specific factors on the clearance rate of circulating cells. By manipulating donor and recipient mice, we can observe how changes to either circulating or non-circulating factors can directly impact the clearance of CLCs. We demonstrate that changes in the bone marrow, such as the presence of significant leukemia engraftment, as well as treatment status of disease can alter the clearance profile of CLCs. Our data also suggests that the adhesion molecule E-selectin plays an important role in modulating the clearance rates of CLCs. E-selectin has been explored as a potential therapeutic target for concurrent chemotherapy in AML^23^, and the data we have shown here supports this and suggests that other types of leukemia, such as ALL, may also benefit from E-selectin based therapies.

Two core findings of this work are that the circulation time of leukemias may be substantially longer than that found in circulating solid tumor cells and that this circulation time is impacted by the presence of existing disease at sites of leukemia dissemination. While it is perhaps intuitive that blood cancers would have greater propensity to persist in the blood compared to solid tumors, this work highlights that not all disseminating malignancies have acquired similar persistence characteristics. Additionally, this work shows that leukemia cells do not persist for indefinite periods in circulation and that the circulating pool is continually fed from extravascular sites. Thus, this kind of blood exchange system can yield fundamental insight into the dissemination of hematopoietic disease.

That the presence of a significant existing disease burden impacts tumor circulation kinetics is perhaps also not surprising. However, these data have important implications. First, it suggests that dissemination of mutant clones within a leukemia may be impaired in the presence of substantial disease, and that broad dissemination of mutant clones may be facilitated by the ablation of bulk disease. Second, our data shows that eradication of tumor burden is not the same as eradication of normal bone marrow resident cells in promoting tumor extravasation and engraftment. How conditioning agents used to prepare patients for stem cell transplantation or adoptive cell therapy impact donor cell engraftment is a fundamental question – one that may be uniquely informed using this type of blood exchange approach.

Our studies also demonstrate that leukemias may experience profound changes in circulation profiles after treatment. While the idea that leukemia adhesion can mediate drug resistance is well-established, the notion that changes in cell adhesiveness and circulation kinetics can stably persist in relapsed tumor cell populations is less appreciated. Indeed, it is logical and concordant with our findings that relapsed tumor cells may survive both initial and subsequent therapies due to their greater propensity to avoid circulation and reside in extravascular sites.

Finally, our findings emphasize a role for adhesion proteins as mediators of therapeutic resistance and potential targets. Due to the discontinuous basement membrane in the sinusoidal capillaries of the bone marrow ^44,45^, E-selectin expressed on BMECs is directly exposed to both the blood and marrow space^46–48^. In the setting of leukemia, high levels of E-selectin on BMECs likely bind leukemia cells both along the vascular walls and in the marrow, creating a static environment with minimal mixing between the blood and bone marrow compartments, thereby inhibiting leukemia cells from exiting circulation. By blocking E-selectin in this environment, leukemia cells in the bone marrow could be released and more free to interchange between tumor compartments. This hypothesis agrees with our observation that infused fluorescent CLCs can clear more rapidly in tumor-bearing recipients after treatment with an E-selectin antibody. Given recent results that uprolesolan, a small molecule E-selectin inhibitor, can increase the potency of chemotherapy in AML^22,23^, a central question is whether its efficacy is predominately achieved through local interactions in the marrow, or due to alterations in CLC dynamics. Our system is well positioned to address this type of question and better understand how novel drugs can impact the circulation kinetics of CLCs, which may change the interpretation of how blood burden in the patient relates to therapy efficacy.

Our study has multiple limitations, including a lack of validation in other mouse models, such as patient-derived xenograft (PDX) models, as well as a lack of genetic perturbations, such as knock-out models of adhesion proteins. However, the goal of this study was to validate the use of this technology to investigate biological questions. Future studies will explore whether these findings are generalizable across models and identify the mechanism by which specific factors can contribute to alterations in circulation properties of CLCs.

Because our system can decouple the influence of circulating and non-circulating factors, we envision that this blood exchange system can be broadly applied across different cancer types to understand the clearance rates of circulating cells and properties that govern cell clearance, from homing and adhesion factors to mechanisms of drug resistance.

## Methods

### Blood exchange system

The platform used involves the connection of two Mouse CTC Sorter systems^11,12^. Each system consists of a microfluidic device, a laser based optical detection system, and a computer controller. The PDMS microfluidic chip is a 100µm wide x 45µm tall channel of 1cm in length, with a single inlet and a single outlet bonded to a glass slide. The optical detection system consists of a 532nm laser that is split into two using polarizing beam splitters and is focused into lines on the microfluidic channel using a cylindrical lens. Emitted light passes through the dichroic and is detected by a photomultiplier tube (PMT). The computer controller records the PMT signal and identifies the presence of fluorescent CLCs by their signature “double-peak” as they pass through the two laser lines. The computer also controls the peristaltic pump which pushes blood through the connective tubing and microfluidic chip at a set rate of 60µL/min, and the two cameras which observe two regions of the channel to monitor the system.

### Model fitting

To identify the concentration of CLCs in the blood, the raw data from the PMT is fed into a MatLab program to identify the peak signatures, and the Equations 1 & 2 described in the Results section are used to identify the circulation parameters. For Equation 2, while the value of “k” is the variable of interest in the equation, the variables “a” and “c” are scaling factors used solely to find the best fit for “k”. The variable “c” describes the steady state level of circulation in the mice but is much less robust of a measurement than the fraction remaining value. Especially when there is a high concentration in the blood at the end of blood exchange, small changes in measurements in the last few minutes can drastically impact the best-fit “c” value, with relatively little impact on the “k” equilibration rate. Similarly, the variable “a” describes the amplitude from the estimated “c” to the starting concentration, and thus can also fluctuate with small changes in measurements. As such, the scaling factors “a” and “c” are not used for downstream analysis.

### Mice

All animal procedures were approved by the Massachusetts Institute of Technology Committee on Animal Care (CAC), Division of Comparative Medicine (DCM). Animals were housed on hardwood chip bedding, with a 12/12 hour light-dark cycle at a temperature of 70°F +/-2 and humidity of 30-70%. All mice used for these studies were C57BL/6 mice procured from Jackson Laboratories.

Because the cancer cell lines used were derived from male mice, all mice in these studies were male. Tumors were initiated when the mice were aged 12-16 weeks. Cells were thawed 1-3 days prior to tumor initiation and passaged daily. On the day of tumor initiation, cells were resuspended in sterile PBS and 500k cells in 100uL were injected via tail vein into each mouse. For the AML model, 1x 5Gy of radiation was dosed 24 hours before injection of tumor cells to allow for tumor engraftment.

To access the circulation for blood exchange experiments, surgical cannulation was performed. Catheters (Instech: C20PU-MJV1458 and C10PU-MCA1459) were placed into the carotid artery and jugular vein of the mice and were externalized through a vascular access button (Instech: VABM2JB/25R25). Catheters were flushed every 3-4 days with saline before filling with a heparinized locking solution (SAI HGS-10-C). Blood exchanges for untreated mice were performed 6-19 days post tumor initiation.

Each blood exchange is performed with a new donor and recipient mouse.

### Drug testing

For the ALL model, a single 50mg/kg dose of the chemotherapy cyclophosphamide (CTX) was given intraperitoneally (IP) 8 days after tumor initiation of 500k cells via tail vein injection. We found that the CTX treated mice had a 2–3-week life extension compared to the untreated animals (Sup. 10a). For the AML model, we used a 5+3 treatment of cytarabine (araC) and doxorubicin (dox). For 5 days, beginning 8 days post tumor initiation of 500k AML cells, the mice were dosed IP with 20mg/kg araC. And on the first three of those days (days 8-10 post tumor initiation), mice were given a concurrent dose of 2mg/kg dox. The 5+3 araC/dox treatment provided roughly one week life extension (Supp. 10b).

Blood exchange with relapse AML mice was performed 9-16 days post final day of treatment, and blood exchange with relapse ALL mice was performed 13-16 days post final day of treatment.

### Flow cytometry

Bone marrow cells were isolated through manual grinding of left and right femurs with mortar and pestle, and red blood cells removed through ACK lysis buffer before 10 minutes FcX blocking BioLegend 101320) and 20 minutes of staining at 4°C. Blood samples were collected from terminal cardiac puncture, and red blood cells were removed with two rounds of ACK lysis buffer before 10 minutes FcX blocking and 20 minutes of staining at 4°C. Flow cytometry was performed on a BD LSR HTS-2 analyzer. The gating strategy used to identify the fraction of BMECs expressing adhesion proteins is shown in Supp. 11. After selecting for single cells through forward and side scatter, endothelial cells were selected as DAPI-(live), CD31+ (endothelial marker [BioLegend: 102410]), and CD45-(white blood cell marker [BioLegend: 103108). A cutoff on the endothelial cells in the channel of the adhesion molecule antibody (E-selectin [Santa Cruz Biotechnology: sc-59766 PE] and VCAM1 [Life Technologies: 11-1061-82 FITC]) was used to identify high expressing cells within the BMECs. Because of the low abundance of endothelial cells in the marrow <1%, at least 1 million cells were analyzed per mouse bone marrow sample. To identify the expression of adhesion markers on the leukemia cells, the following strategy was used. To assess for E-selectin binding potential, a recombinant E-selectin human IgG chimera [BioLegend: 755504] was conjugated to fluorescent α-Human IgG [Life Technologies: A10631] for 30 minutes at room temperature prior to staining^22^. The presence of integrin was determined with an integrin β1 antibody [Life Technologies: 11-0291-82]. After selecting for single cells through forward and side scatter, tumor cells were selected as DAPI-(live), CD45+ (white blood cell marker [BioLegend: 103114]), PE+ (constitutively expressed RFP). Average expression in the channel of the adhesion molecule (E-selectin binding or integrin β1) was then used to assess binding potential.

### E-selectin interference

To interfere with BMEC expression of E-selectin, 100µg (100µL) of mouse E-selectin antibody (BioLegend: 148803) was injected retro-orbitally 20 minutes prior to blood exchange in the recipient mouse. To interfere with the expression of E-selectin binding molecules on tumor cells, 20µg (100µL) of recombinant mouse E-selectin (BioLegend: 755504) was injected retro-orbitally 20 minutes prior to blood exchange.

### Histology

Femurs were isolated and fixed in formalin for 24 hours before being washed and stored in 70% ethanol. Samples were prepared in paraffin wax and sliced into 5µm sections before hematoxylin and eosin staining. Slides were analyzed on a Nikon A1R confocal microscope.

## Data availability

The data that support the findings of this study are available from the corresponding authors upon reasonable request.

## Acknowledgments

We thank the Koch Institute’s Robert A. Swanson (1969) Biotechnology Center for technical support, specifically the Flow Cytometry Core Facility, the Hope Tang (1983) Histology Facility, and the Microscopy Core Facility. We thank the MIT Division of Comparative Medicine for support and guidance of animal experiments. This work was supported in part by the Virginia and D.K. Ludwig Fund for Cancer Research and Stand Up to Cancer (SU2C) Convergence Program 3.1416. D.M.W. was supported by NCI R35 CA239158 and NCI P01 CA248384.

## Contributions

A.B.M, D.M.W., M.T.H., and S.R.M. conceptualized the study. A.B.M., A.L., F.H.R., D.G., A.A.L., D.M.W., M.T.H., and S.R.M. designed the experiments. A.B.M, A.L., F.H.R., L.L., and C.B., performed the blood exchange experiments. L.L. performed the animal surgeries. A.B.M. analyzed blood exchange data. A.B.M. and A.L. performed flow cytometry. A.B.M., A.L., S.D., and Y.Z. measured the leukemia cell biophysical properties. A.B.M., M.T.H., and S.R.M. wrote the paper with input from all the other authors.

