## Supplemental Figures for "Leukemia circulation kinetics revealed through blood exchange method"

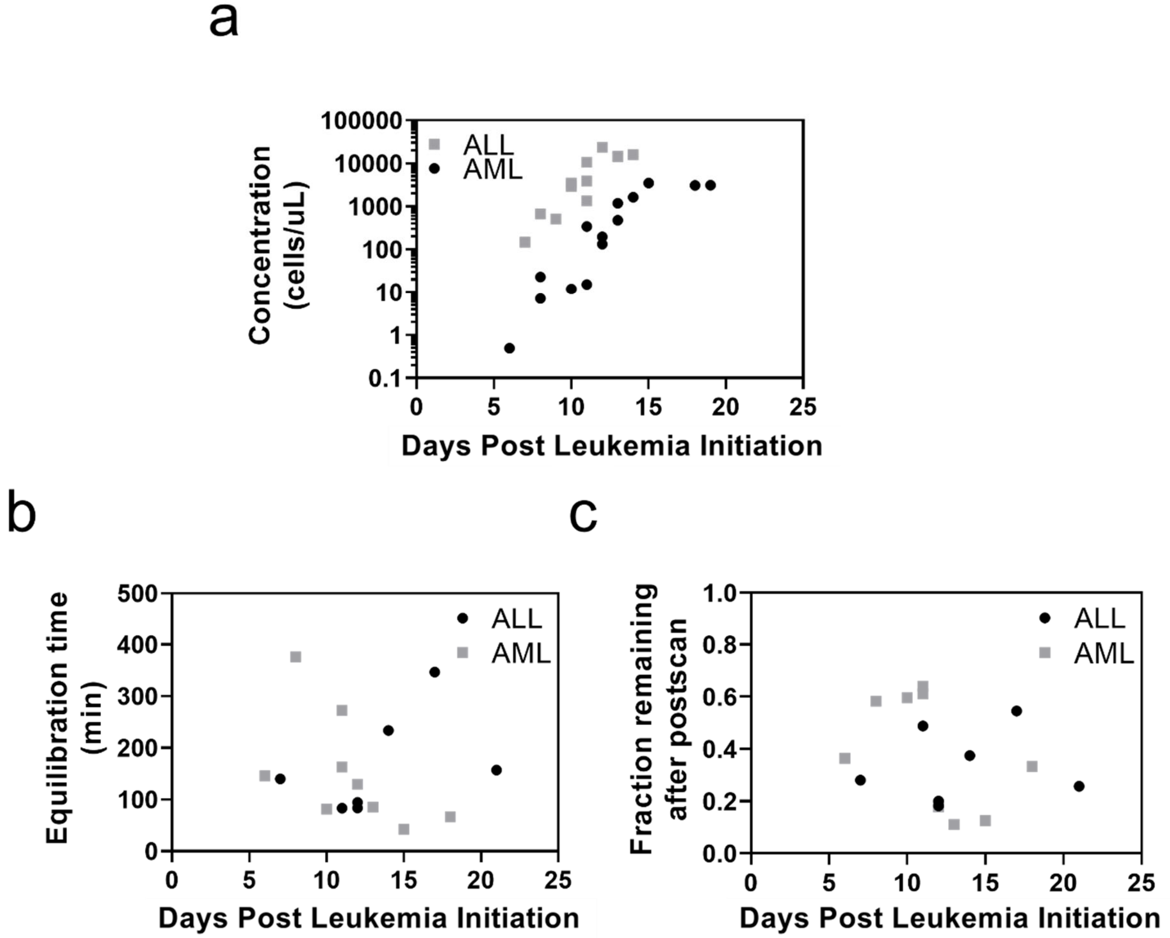


**Supplemental Figure 1 | No changes in circulation kinetics observed in response to tumor burden.** (a) Strong correlation in both AML ($R^2$=0.7163, p=0.0001) and ALL ($R^2$=0.5879, p=0.0036) between days post tumor initiation and circulating concentration of CLCs using an exponential growth fit. (b,c) No significant linear correlation between days post tumor initiation and either circulation kinetic, equilibration time (AML: $R^2$=0.3219, p=0.1111; ALL: $R^2$=0.2104, p=0.3005) or fraction remaining at the end of the post-blood exchange scan (AML: $R^2$=0.2119, p=0.2125; ALL: $R^2$=0.0230, p=0.7458).

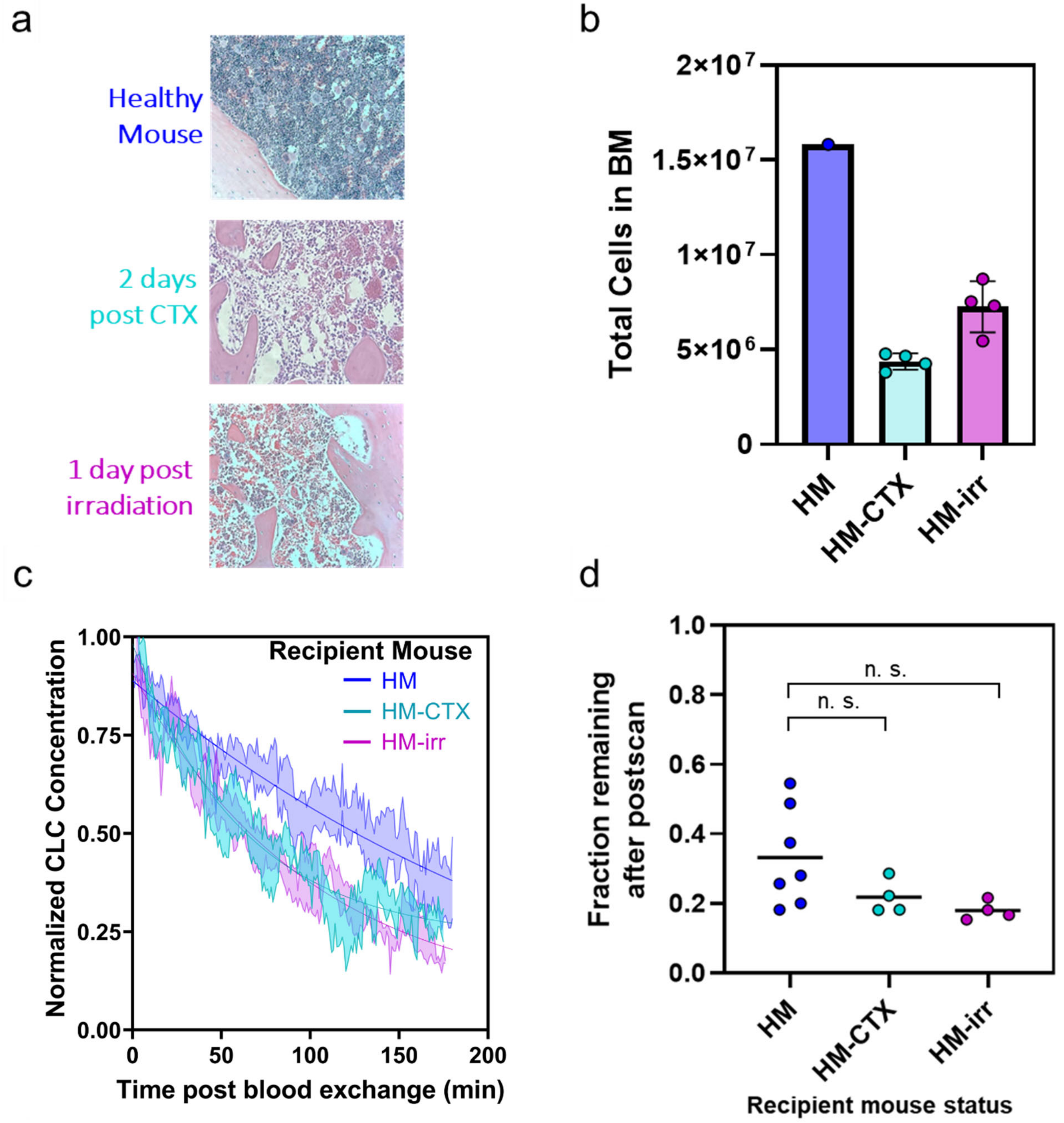


**Supplemental Figure 2 | Hypocellularity is insufficient to impact circulation kinetics.** (a, b) Healthy mice (HM) treated with cyclophosphamide (CTX) or irradiation (1x 5Gy) show 50-70% reduction in total cells in the marrow. (c, d) Clearance of RFP+ ALL donor CLCs is not significantly impacted by hypocellularity as measured by fraction remaining (Tukey multiple comparisons p=0.2237 and 0.0868 for HM-CTX and HM-irr, respectively). For decay profiles (c), shaded regions are represented by mean +/- standard error, and lines represent the best fit decay curve. Each dot represents an independent donor-recipient mouse pair.

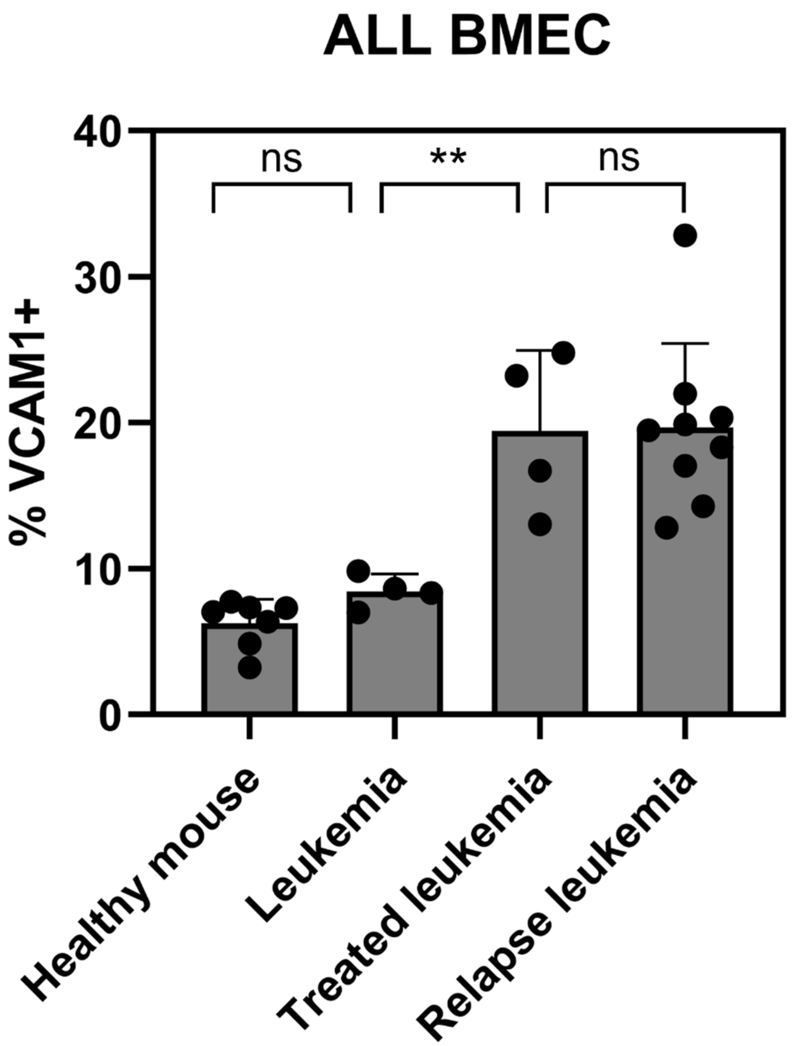


**Supplemental Figure 3 | VCAM1 expression on bone marrow endothelial cells (BMEC) in ALL does not correlate with disease status.** Percent of BMECs with positive expression of VCAM1 throughout tumor and treatment in ALL. Bars represent mean +/- standard deviation. Tukey multiple comparisons test **p=0.0093

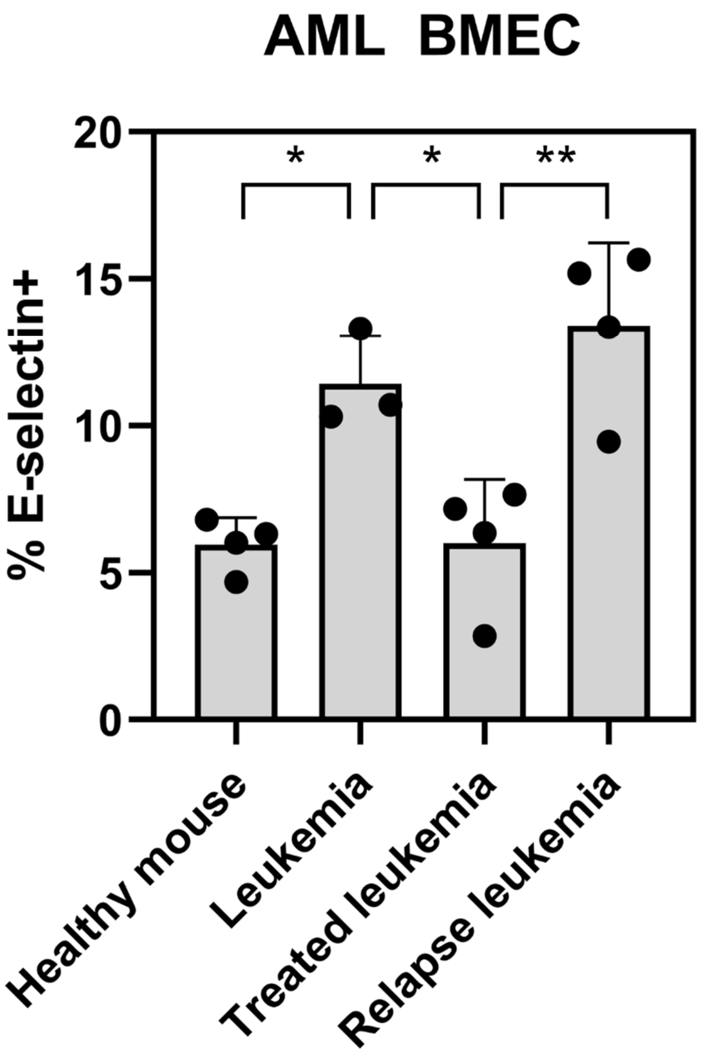


**Supplemental Figure 4 | E-selectin expression on bone marrow endothelial cells (BMEC) in AML increases in diseased context.** Percent of BMECs with positive expression of E-selectin throughout tumor and treatment in AML. Bars represent mean +/- standard deviation. Tukey multiple comparison test p=0.0217, 0.0227, and 0.0016 respectively from left to right.

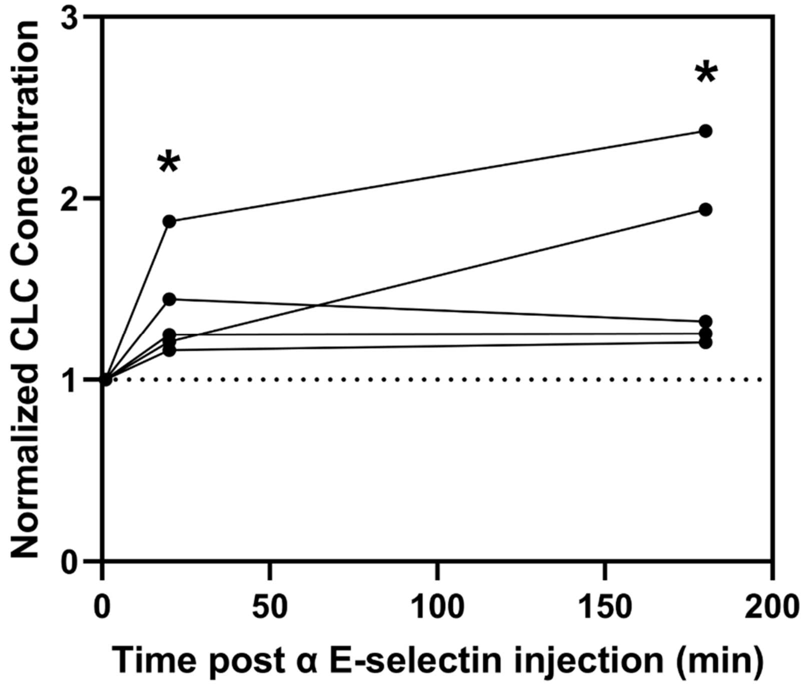


**Supplemental Figure 5 | E-selectin antibody injected intravenously can recruit cells into circulation in as short as 20 minutes.** Change in circulating concentration of CLCs after injection of E-selectin antibody into leukemia bearing mice. Two tailed t-test relative to baseline- 20 min: $p=0.0180$, 3 hour: $p=0.0276$.

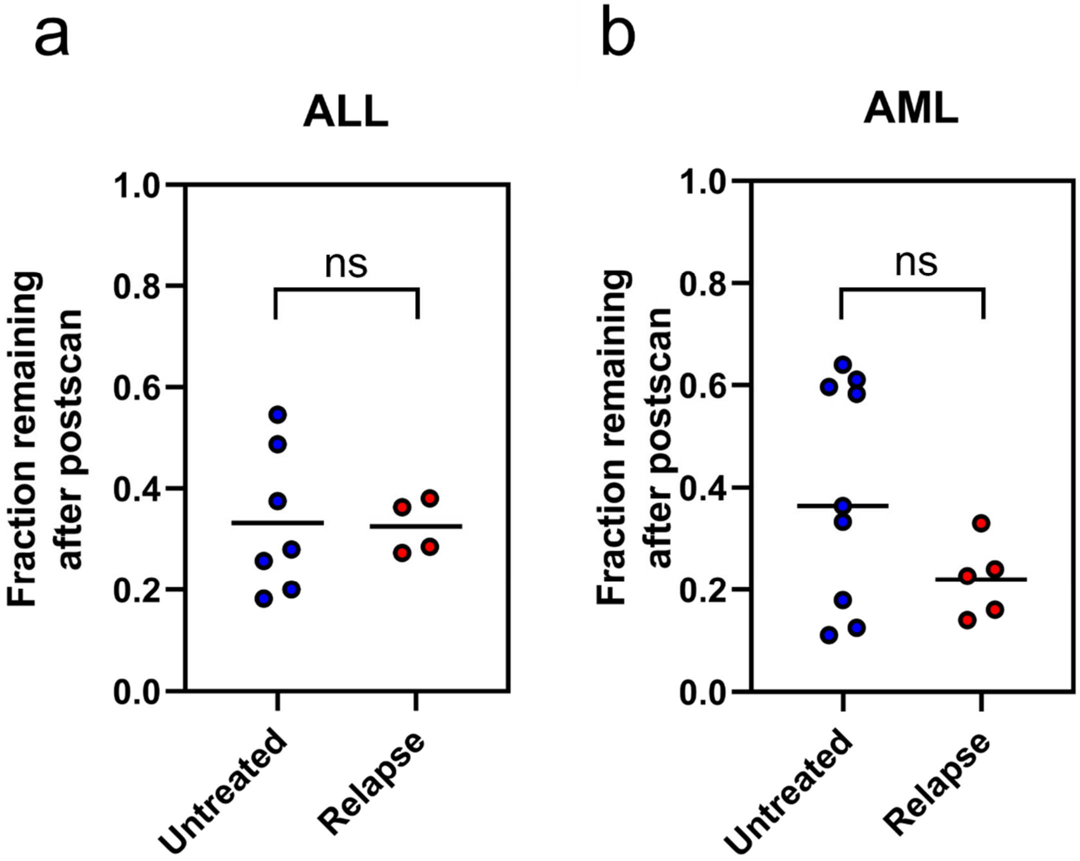


**Supplemental Figure 6 | Fraction remaining of untreated and relapse leukemia in healthy recipients.** No significant change observed in fraction remaining at relapse in (a) ALL (two-tailed t test p=0.9295) or (b) AML (two-tailed t test p=0.1169) models.

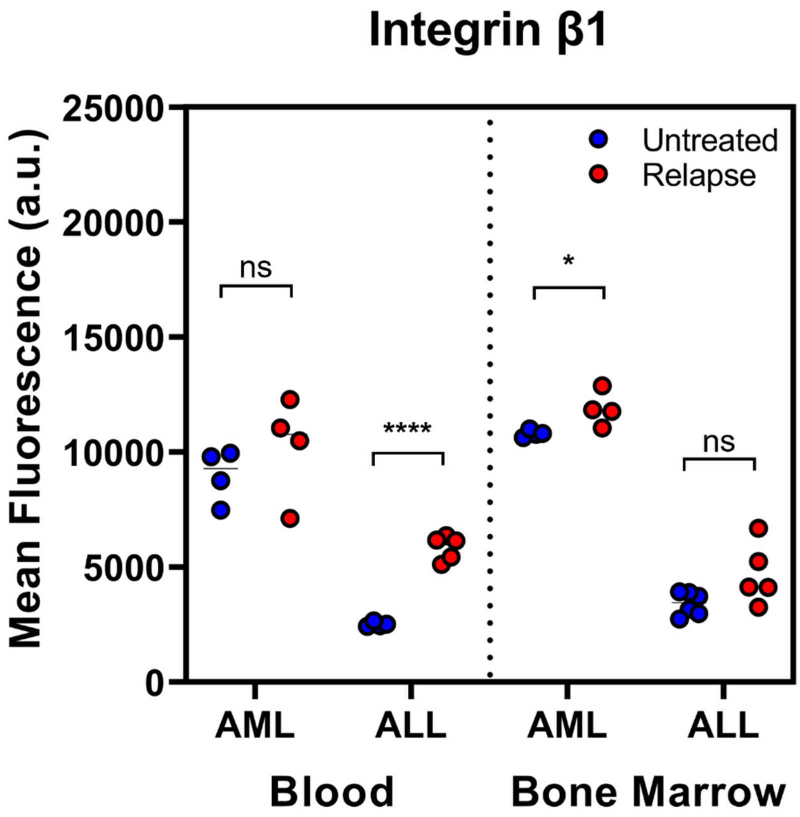


**Supplemental Figure 7 | Integrin β1 expression on leukemia cells.** Some tumor compartments showed increases in expression of integrin β1 on leukemia cells (two tailed t-tests: *p=0.0317, ****p<0.0001).

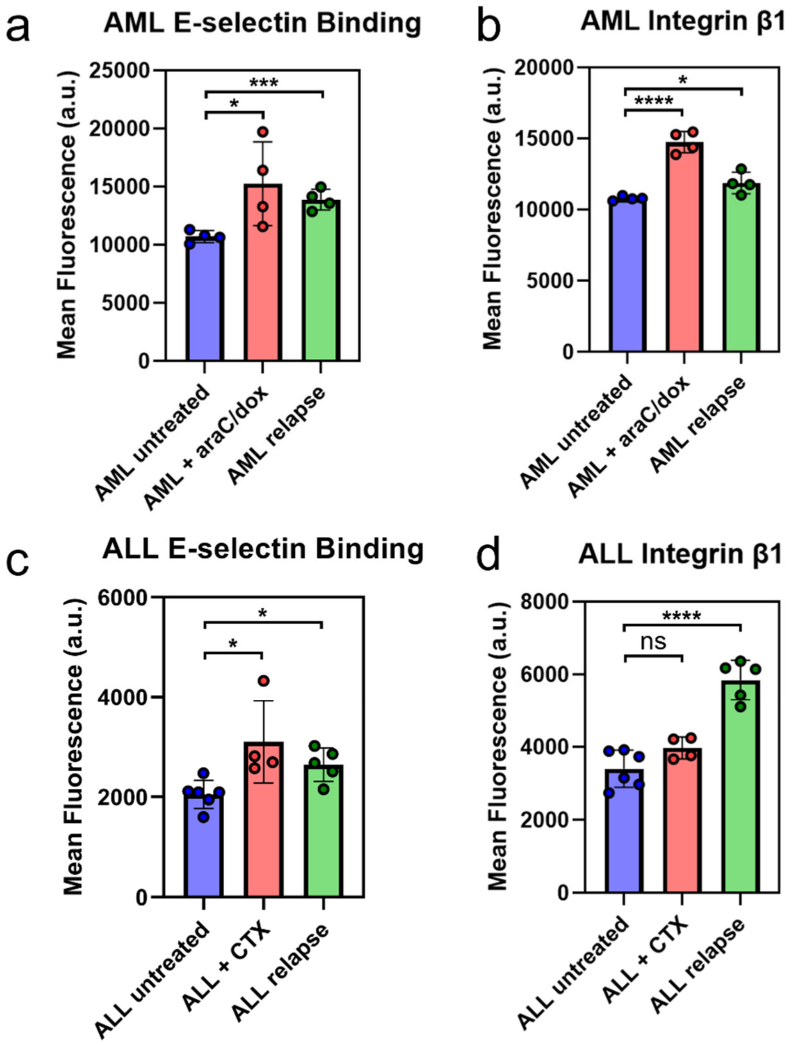


**Supplemental Figure 8 | Leukemia surface adhesion molecules increase on leukemia cells of the bone marrow upon chemo treatment and remain elevated through relapse.** (a,b) AML and (c,d) ALL leukemia cells show an increase in expression of adhesion molecules after treatment and at relapse. (Two-tailed t tests: *p<0.05; ***p<0.001; ****p<0.001).

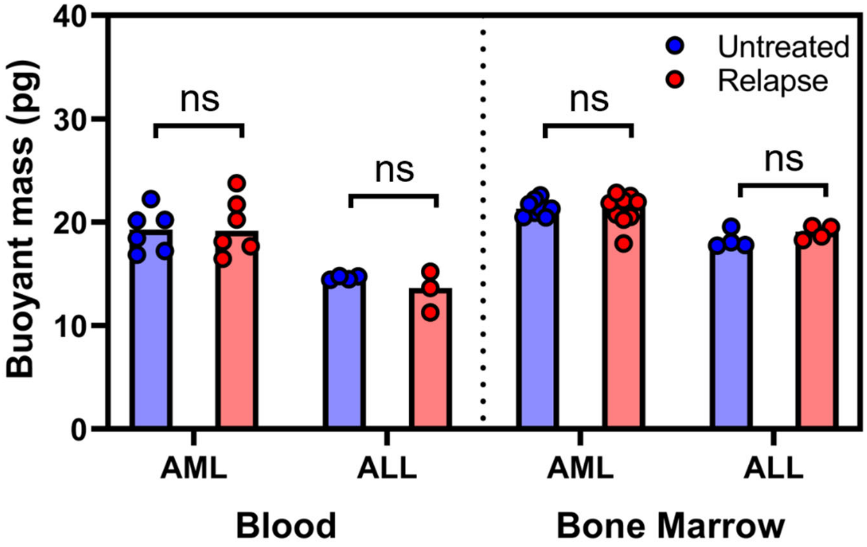


**Supplemental Figure 9 | Buoyant mass measurements of leukemia cells.** No change in buoyant mass seen using suspended microchannel resonator to observe single cell buoyant mass. Each dot indicates the mean mass for a given mouse sample, with >200 single cell measurements per sample.

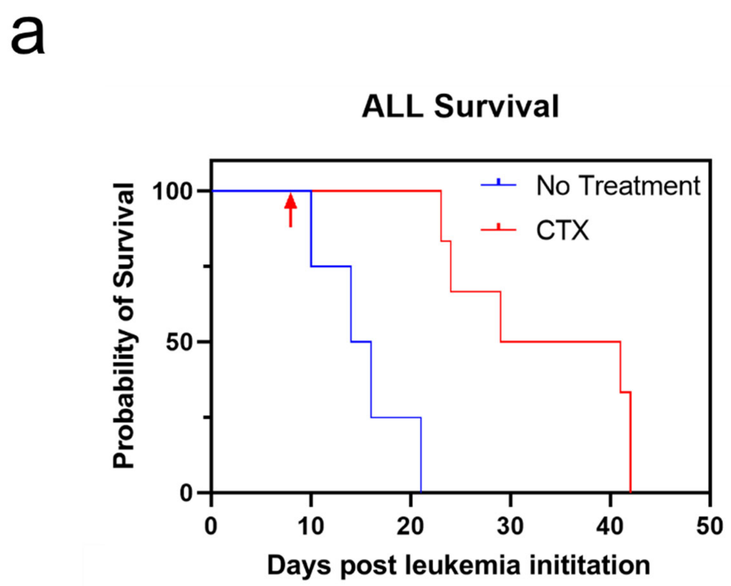


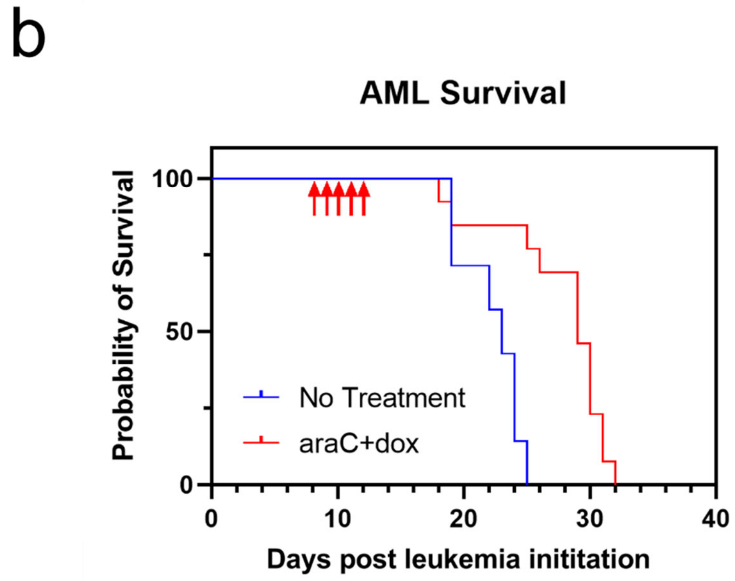


**Supplemental Figure 10 | Chemotherapy treatment for leukemia.** (a) 1 dose of cyclophosphamide (CTX) was given 8 days post treatment to induce a 2–3-week extension of life (Mantel-Cox test p=0.0011). (b) A 5+3 regimen of cytarabine (araC) and doxorubicin (dox) was administered beginning on day 3. For the first three days of treatment, both drugs were given, and on days 4 and 5, only araC was given. This induced approximately a 1-week extension of life (Mantel-Cox test p=0.0008).

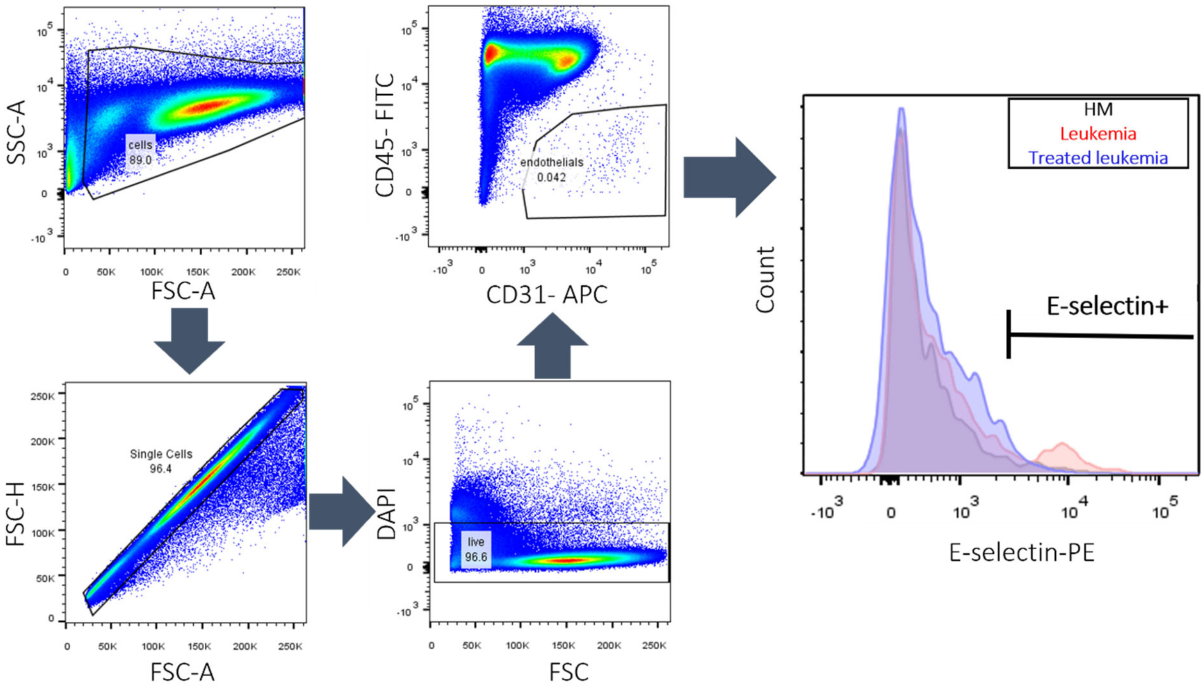


**Supplemental Figure 11 | Gating strategy for determining E-selectin expression of BMECs.** After gating out single cells, live (DAPI-) cells were gated. Endothelial cells were identified as CD31+/CD45-, and E-selectin+ BMECs were gated to determine %E-selectin+. Because endothelial cells were very rare, at least 1 million events were collected for analysis.
